# A New Assay Capturing Chromosome Fusions Shows a Protection Trade-off at Telomeres and NHEJ Vulnerability to Low Density Ionising Radiation

**DOI:** 10.1101/2021.05.04.442438

**Authors:** Sabrina Pobiega, Olivier Alibert, Stéphane Marcand

## Abstract

Chromosome fusions threaten genome integrity and promote cancer by engaging catastrophic mutational processes, namely chromosome breakage-fusion-bridge cycles and chromothripsis. Chromosome fusions are frequent in cells incurring telomere dysfunctions or those exposed to DNA breakage. Their occurrence and therefore their contribution to genome instability in unchallenged cells is unknown. To address this issue, we constructed a genetic assay able to capture and quantify rare chromosome fusions in budding yeast. This chromosome fusion capture assay (CFC) relies on the controlled inactivation of one centromere to rescue unstable dicentric chromosome fusions. It is sensitive enough to quantify the basal rate of end-to-end chromosome fusions occurring in wild-type cells. These fusions depend on canonical nonhomologous end-joining (NHEJ). Our results show that chromosome end protection results from a trade-off at telomeres between positive effectors (Rif2, Sir4, telomerase) and a negative effector partially antagonizing them (Rif1). The CFC assay also captures NHEJ-dependent chromosome fusions induced by ionising radiation. It provides evidence for chromosomal rearrangements stemming from a single photon-matter interaction.

## Introduction

Chromosome stability requires that DNA double-strand breaks (DSBs) are repaired by canonical NHEJ or homologous recombination. NHEJ is an efficient pathway directly resealing DSB ends, most often accurately (1–4). In line with other DNA repair pathways, NHEJ repair sometimes comes with errors. Its contribution to mutagenesis is twofold: sequence changes at the break sites and chromosomal rearrangements (1, 2, 5–10). NHEJ-dependent chromosomal rearrangements include dicentric chromosomes that lead to catastrophic mutational processes, namely chromosome breakage-fusion-bridge cycles and chromothripsis (11–13). The rarity of these events is key to genome integrity.

NHEJ’s Achilles heel is its susceptibility to the co-occurrence within the cell nucleus of multiple DSBs, a situation that ionising radiation easily create (14–17). This co-occurrence favours DNA end erroneous resealing stemming from distinct breaks, resulting in chromosome translocations. NHEJ error frequency results from a balance between broken end diffusion and their synapsis (tethering) by the NHEJ machinery (5, 9, 18, 19).

An efficient NHEJ machinery must also co-exist in the nucleus with stable telomeres, the native chromosome ends. This restriction to DSB is an important element of NHEJ accuracy. It is based on a strong inhibition of NHEJ at telomere ends which ensures that chromosome ends rarely fuse. NHEJ inhibition relies on a limited number of proteins present at telomeres. Defects in these proteins or telomere shortening, which reduce their amount, compromise NHEJ inhibition and lead to frequent chromosome fusions (12, 20–24). Chromosome fusions may occur in unchallenged cells, but due to technical limitations their frequency and therefore the absolute efficiency of NHEJ inhibition at functional telomeres remains difficult to assess.

The rate of telomere fusions has been explored in the *Schizosaccharomyces pombe* and *Saccharomyces cerevisiae* yeast model organisms. In fission yeast, a genetic assay capturing fusions between the telomeres of a plasmid-based mini-chromosome shows that telomeres in unperturbed wild-type cells may be more prone to NHEJ-dependent fusions than initially thought (approximately 10^−4^ events per cell), with the caveat that mini-chromosome telomeres may not be fully functional (25). In budding yeast, a genetic assay capturing NHEJ-dependent fusions between an endonuclease-induced broken end and native telomeres shows that this type of events is rare in wild-type cells, with a frequency around 10^−7^ fusions per cell (26). This sensitive assay misses the fusions between native telomeres. PCR-based assays that detect the frequent NHEJ-dependent fusions occurring between dysfunctional telomeres in mutant contexts fail to capture the rarer fusions that may occur in wild-type cells (27–32). The detection threshold of these PCR-based assays is approximately 10^−5^-10^−6^ fusions per cell. Therefore, a quantitative knowledge of telomere fusion contribution to genome instability in normal cells is still missing.

This led us to construct a genetic assay able to capture and quantify rare chromosome fusions in *S. cerevisiae*. This assay relies on the controlled inactivation of one centromere to rescue unstable dicentric chromosome fusions. Sensitive enough to explore wild type contexts, it shows the rarity of telomere fusions in wild-type yeast cells (less than 10^−6^ events per cell) and the synergy of the pathways inhibiting NHEJ at telomeres. Telomere length is also a key determinant of telomere protection efficiency since telomerase loss causes a rapid increase in fusion frequency. Unexpectedly, this new assay also captures NHEJ-dependent chromosome fusions induced by ionising radiation. Our data suggest that a single photon-matter interaction can be sufficient to generate an NHEJ-dependent rearrangement.

## Results

### A new assay capturing rare NHEJ-dependent chromosome rearrangements

Chromosome fusions create unstable dicentric chromosomes. The inactivation of one centromere is an efficient way to stabilize dicentric chromosomes and to select chromosome fusions (33–35) (**Fig. 1A**). In a previous study, we inactivated the native centromere of *S. cerevisiae* chromosome 6 (*CEN6*) using inducible promoters pointed toward the centromere (34). However, the assay lacks sensitivity to capture fusions in wild-type cells (34). To circumvent this limitation, we created an assay where the Cre recombinase eliminates the centromere. To increase the assay specificity, a promoter and a *LEU2* coding sequence surround a *loxP* cassette that includes *CEN6* and a *URA3* gene (**Fig. 1B**). Cre expression from a galactose-inducible promoter simultaneously deletes *CEN6* and generates a functional *LEU2* gene. The latter allows a counterselection of cells having failed Cre-induced recombination (approximately 1 every 10^5^ cells) by screening for leucine prototrophic *cen6-Δ* clones on galactose medium lacking leucine. Note that we chose chromosome 6 because the mis-segregation of this chromosome leads to two unviable cells, increasing the stringency of the assay (34, 36).

**Figure 1.**
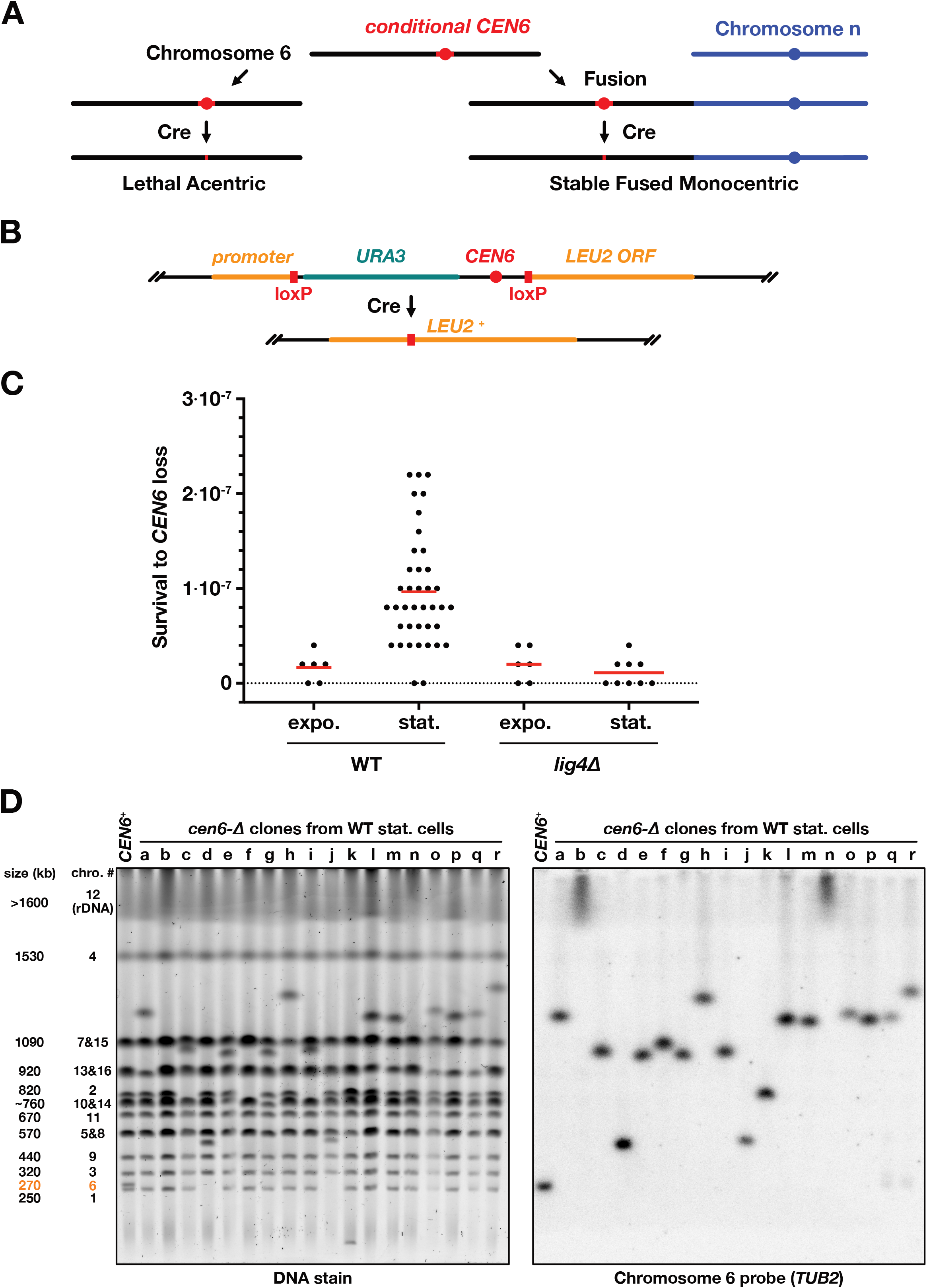
An assay to capture chromosome fusions. (**A**) *CEN6* elimination stabilises fusions between chromosome 6 and another chromosome. If chromosome 6 is unfused, *CEN6* elimination generates an unstable acentric chromosome leading to cell death. (**B**) Schematic representation of the loxP cassette inserted at *CEN6*. (**C**) Survival to *CEN6* loss of wild-type and NHEJ-deficient (*lig4Δ*) cells. Expo.: cells growing exponentially in rich medium (OD_600nm_ <1). Stat.: cells reaching stationary phase in rich medium (3 days, (OD_600nm_ ≈25). (**D**) Karyotype of *cen6Δ* clones from wild-type stationary cells. In clones *b* and *n*, chromosome 6 is fused to chromosome 12.

First, we estimated the frequency of survival to *CEN6* loss in wild-type cells (**Fig. 1C**). Cells growing exponentially (expo.) or having reached stationary phase (stat.) in liquid glucose rich medium were spread on galactose medium plates lacking leucine (5 × 10^7^ cells per plate; each data point from a distinct cell culture). Leucine-prototrophic colonies emerge after 4-5 days at 30°C. Loss of the *URA3* gene and PCR on a sample of survivors confirm *CEN6* loss. From exponentially growing cells, the frequency of survivors to *CEN6* loss is low, about 2 × 10^−8^ events per cell (**Fig. 1C**, 0 to 2 colonies per plate, close to the practical limit of the assay sensitivity). This frequency increases about 5-fold to 10^−7^ events per cell when cells have reached stationary phase prior to plating. This shows that events leading to survival accumulate in stationary cells, either because they stop being counter-selected in non-dividing cells or because they form more frequently in those cells or both. The increase in stationary cells also indicates that most selected events form prior to plating and centromere elimination. By contrast, rarer events selected from exponentially growing cells might also be secondary outcomes of centromere elimination (35). Lig4^Dnl4^ is the DNA ligase essential to canonical NHEJ (37). Remarkably, in the absence of Lig4, survival remains low when stationary cells are plated on galactose medium lacking leucine (≤2 × 10^−8^ events per cell, **Fig. 1C**). This shows that the events leading to survival to *CEN6* loss and increasing in stationary wild-type cells are mostly products of canonical NHEJ.

To determine the chromosome rearrangements captured by the assay, we analysed by pulse-field gel electrophoresis the karyotype of individual *cen6-Δ* clones obtained from stationary wild-type cells. **Figure 1D** shows a representative sample of selected clones. In all clones, chromosome 6 as well as another chromosome (**Fig. 1D** left panel) are missing at their native position. In most cases, a single new chromosome appears at a size close to the sum of the two missing chromosomes, as expected for chromosome fusions. The fused chromosome hybridizes with a probe from chromosome 6 left arm (**Fig. 1D** right panel). Thus, the assay can capture individual chromosome fusions. In a few clones, two new chromosomes appear, suggestive of a reciprocal translocation (e.g. clone *k* in **Fig. 1D**, the smaller of the rearranged chromosomes likely an acentric in multiple copies). In conclusion, the assay captures rare NHEJ-dependent chromosome fusions occurring in stationary wild-type cells.

### Telomere fusions in wild-type cells

Chromosome fusions can stem from fusions between telomeres or be consecutive to telomere losses or internal breaks. To distinguish between these events, we used a Southern blot approach to first determine which chromosome end of chromosome 6 is fused. The sequence adjacent to the right end telomere of chromosome 6 (*TEL6R*) is unique. As shown in **Figure 2A** (left panel, same clones as in Fig. 1D), instead of the characteristic smear of intact telomeres, some captured chromosomes fusions display a discreet-size band, indicating that *TEL6R* is fused (e.g. clones *a*, *l* and *n*), or no signal at *TEL6R* (e.g. clone *d*), indicating that this telomere is lost. Among the 47 clones with a chromosome fusion that were tested at *TEL6R*, 11 are fused at *TEL6R* (~23%) and 3 have lost *TEL6R* (~6%).

**Figure 2.**
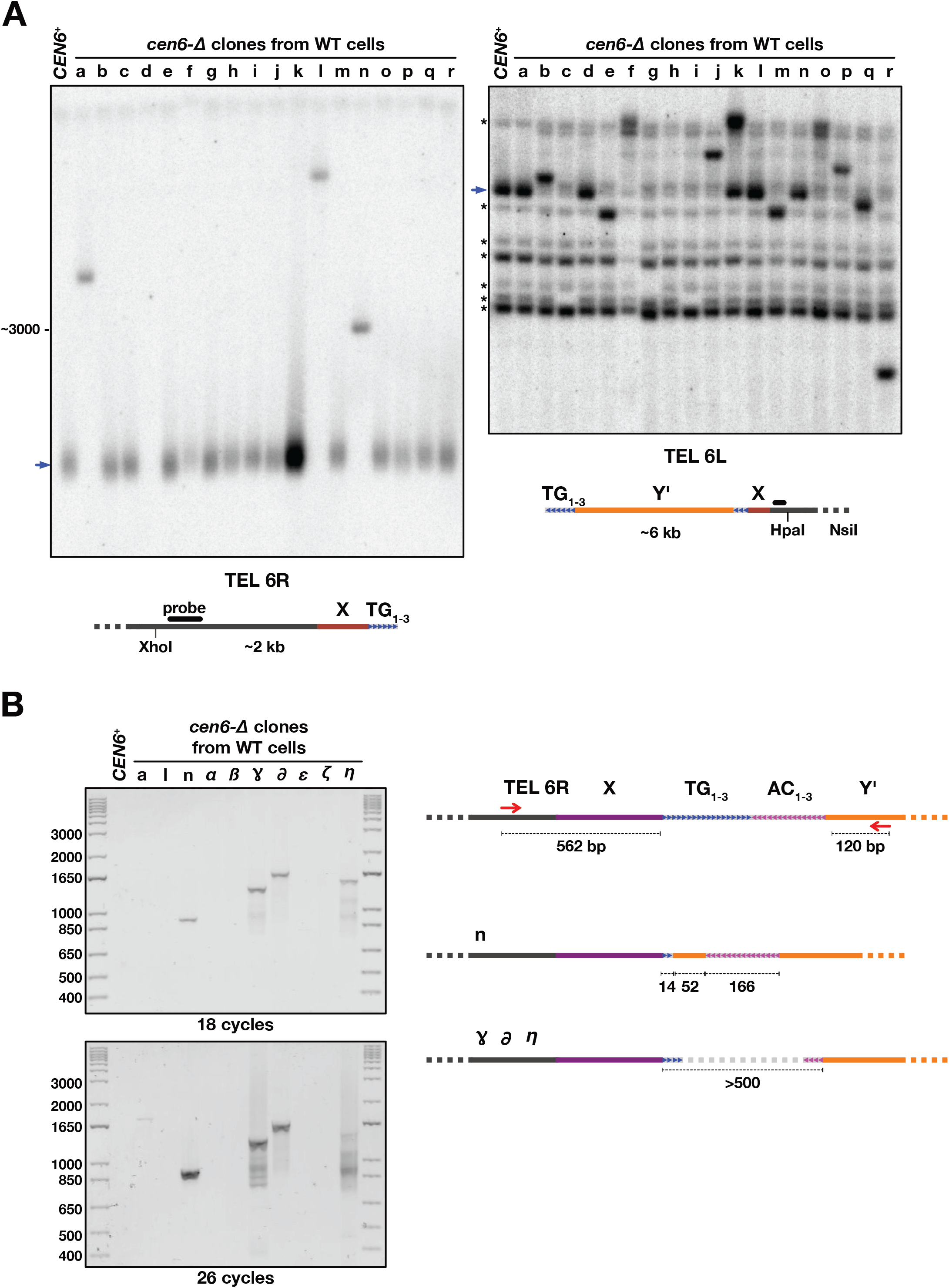
Telomere fusions in wild-type cells. (**A**) Telomere fusions at chromosome 6 telomeres among *cen6Δ* clones from wild-type stationary cells. Same clones as in **Fig. 1D**. *TEL6R* and *TEL6L* positions marked with blue arrows. Cross-hybridizing restriction fragments marked with asterisks. (**B**) PCR amplification of fusions between chromosome 6 right telomere (*TEL6R*) and Y’-containing telomeres.

The sequence adjacent to the left end of chromosome 6 (*TEL6L*) is one of the Y’ repeated elements, 5-7 kb sequences present at the end of approximately half of the *S. cerevisiae* telomeres. The first stretch of non-repeated sequences is about 6 kb from the chromosome end and we used this sequence as a probe to identify which clones have fused at *TEL6L* (**Fig. 2A**, right panel; *TEL6L* position marked with a blue arrow and other cross-hybridizing restriction fragments marked with asterisks). As expected, the clones with a rearranged *TEL6R* display an intact *TEL6L* (e.g. clones *a*, *d*, *l* and *n*). Among the other clones, some display a new band indicative of a fusion event preserving at least a part of *TEL6L* (e.g. clones *b*, *e*, f, *j*, *m*, *p*, *q* and *r*). A shorter restriction fragment indicates that the terminal TG_1-3_ telomere sequences from *TEL6L* are lost in the fusion (e.g. clones *e*, *m*, *q* and *r).* Other clones have lost the probed sequence, indicating a more extensive loss of *6L* chromosome end (e.g. clones *c*, *g*, *h*, and *i*). Clones with an apparent reciprocal translocation maintain both chromosome 6 telomere ends (e.g. clone *k*, whose *TEL6R* end is in multiple copies). Among 47 clones with a chromosome fusion that were tested, 19 are fused at *TEL6L* (~40%) and 14 have lost *TEL6L* (~21%).

Overall, a fraction of the captured chromosome fusions involves the loss of one chromosome 6 telomere (~30%). Are TG_1-3_ telomere repeats still present at the fusion point in the remaining clones? Telomeres and subtelomeres are repeated sequences and their head-to-head fusion generates palindromes difficult to amplify and to sequence. Using a primer specific to *TEL6R* and a primer specific to Y’ element distal end, we could amplify some of the fusions at *TEL6R* (4 out of 11) (**Fig. 2B**). Sequencing shows that three possess TG_1-3_ telomere repeats in head-to-head orientation (clones ɣ, ∂ and η, the palindromic fusion points could not be sequenced) and one a fusion between a very short *TEL6R* and a truncated tandem of Y’ elements (clone n) (**Fig. 2B**). Together, these results show that a fraction of the chromosome fusions captured from unchallenged wild-type cells result from a fusion between two native telomeres.

### NHEJ-dependent chromosome fusions in response to defects in telomere function

Next, we asked if the chromosome fusion capture assay can help to better assess the function of individual telomeric factors in chromosome end protection. In budding yeast, the essential protein Rap1 covers TG_1-3_ telomere repeats. Through direct interactions with its C-terminal domain, Rap1 recruits to telomeres other proteins that contribute to telomere functions and telomere length homeostasis. Previous works showed that the Rap1-interacting factors Rif2 and Sir4 act in synergy to prevent NHEJ-dependent telomere-telomere fusions (30–32, 38). This conclusion was based on a PCR assay that could only detect relatively high frequencies of telomere fusions, for instance the ones occurring in cells lacking both Rif2 and Sir4. The PCR assay failed to assess the impact of single mutants (30–32, 38). To address this issue, we tested the individual contribution of these factors to the low frequency of chromosome fusions observed in wild-type cells using the new chromosome fusion capture assay.

As shown in **Figure 3A**, the loss of Rif2 or Sir4 increases survival to *CEN6* loss about 3-fold and 30-fold respectively. This indicates that chromosome fusions are more frequent in these single mutants, in agreement with a previous observation for Rif2 (26). Analysis of individual *cen6-Δ* clones obtained from Rif2 and Sir4 defective cells shows that each clone stems from a single chromosome fusion between chromosome 6 and another chromosome. In cells lacking Rif2, the fusions occur at one of the two telomeres of chromosome 6 with similar probabilities (**Fig. 3B**, **Supplementary Fig. S1**), suggesting that in wild-type cells the Y’-less *TEL6R* telomere relies slightly more on Rif2 than the Y’-containing *TEL6L* telomere. In cells lacking both Rif2 and Sir4, survival to *CEN6* loss further increases to reach a frequency 4 orders of magnitude above the frequency observed in wild-type cells (**Fig. 3A**). This high survival frequency requires Lig4 indicating that NHEJ produces most of the events leading to survival. Together, these results confirm that Rif2 and Sir4 act in synergy to oppose NHEJ at telomeres. In addition, they show that Rif2 and Sir4 are both essential and non-redundant to bring down NHEJ between telomeres to the very low level observed in normal cells.

**Figure 3.**
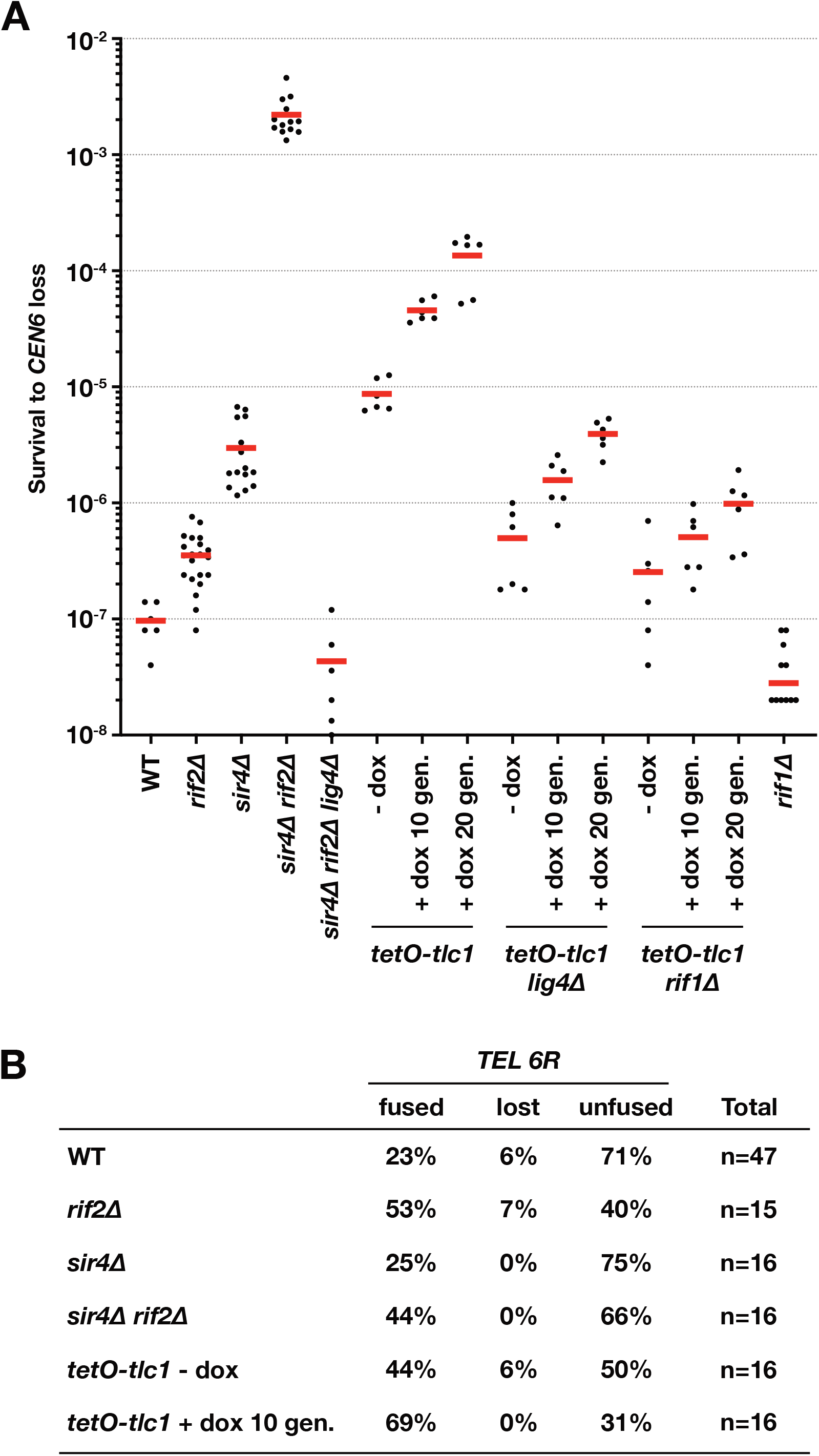
NHEJ-dependent chromosome fusions in response to defects in telomere function. (**A**) Survival frequency to *CEN6* loss in stationary cells lacking Rif2, Sir4, telomerase template RNA (*TLC1*) and Rif1. (**B**) Telomere fusions at chromosome 6 right telomere among *cen6Δ* clones from wild-type and mutant stationary cells.

Next, we addressed the impact of telomere shortening on the occurrence of chromosome fusions. To shorten telomeres, we used a conditional allele of telomerase, *tet0-tlc1*, in which a doxycycline-repressible promoter controls the gene encoding telomerase ARN template *TLC1^TRT^* (39, 40). In uninduced condition (without doxycycline), this allele slightly shortens telomeres (39) (**Supplementary Fig. S2**) and causes a ~100-fold increase in chromosome fusion frequency (**Fig. 3A**). Telomerase inactivation by doxycycline for 10 and 20 population doubling further increases this frequency ~5-fold and 15-fold respectively (~500/1500-fold compared to wild-type cells). These increases require Lig4, indicating that most of the selected events are products of NHEJ. Analysis of individual *cen6-Δ* clones from *tet0-tlc1* cells shows that each clone displays a fusion between chromosome 6 and another chromosome. Fusions at *TEL6R* are more frequent in *tet0-tlc1* cells exposed to doxycycline (**Fig. 3B**, **Supplementary Fig. S1**), suggesting that the Y’ element present at *TEL6L* may offer some buffering in response to a transient telomerase deficiency. In conclusion, these results show that telomerase is essential to chromosome end protection against NHEJ without significant lag at the population level.

We then asked if telomere elongation can compensate for the impact of telomerase deficiency. Rif1 is another telomere factor recruited by Rap1 that represses telomere elongation but does not contribute to telomere protection against NHEJ (30, 41–46). The absence of Rif1 both elongates telomeres and facilitates the binding of Rif2 and Sir4 to telomeres (41, 47, 48). As shown in **Figure 3A**, Rif1 loss strongly reduces the frequency of chromosome fusions induced by the *tet0-tlc1* allele, in both uninduced (− dox) and induced (+ dox) conditions. In addition, the absence of Rif1 alone lowers the basal rate of chromosome fusions of telomerase-proficient cells (~3-fold reduction relative to wild-type). Thus, Rif1 is a moderate but significant impediment to telomere protection against NHEJ-dependent fusions. In conclusion, our data show that telomere protection efficiency in wild-type cells results from a trade-off between positive effectors (Rif2, Sir4, telomerase) and a negative effector partially antagonizing them (Rif1).

### NHEJ-dependent chromosome fusions induced by ionising radiation

Since the assay captures NHEJ-dependent chromosome fusions, we next used it to explore NHEJ repair of co-occurring DNA double-strand breaks (DSBs). To generate random co-occurring DSBs, cells grown to stationary phase in liquid medium were irradiated with gamma rays stemming from ^137^Cs decay (662 keV, 2.60 Gy/min) prior to plating on galactose medium lacking leucine. As shown in **Figure 4A**, irradiation leads to an increase in survival to *CEN6* loss that is correlated to the received dose. The observed inflection at 40 Gy can be explained by an overall loss of cell viability at this dose (**Fig 4B**, **Supplementary Fig. S3A**) (2, 49). The increase in survival to *CEN6* loss requires Lig4 (**Fig. 4A**), indicating that canonical NHEJ generates most of the radiation-induced events selected by the assay. Analysis of individual radiation-induced *cen6-Δ* clones from wild-type cells shows that each clone displays a fusion between chromosome 6 and another chromosome, or, more rarely, a reciprocal translocation (**Fig. 4C**).

**Figure 4.**
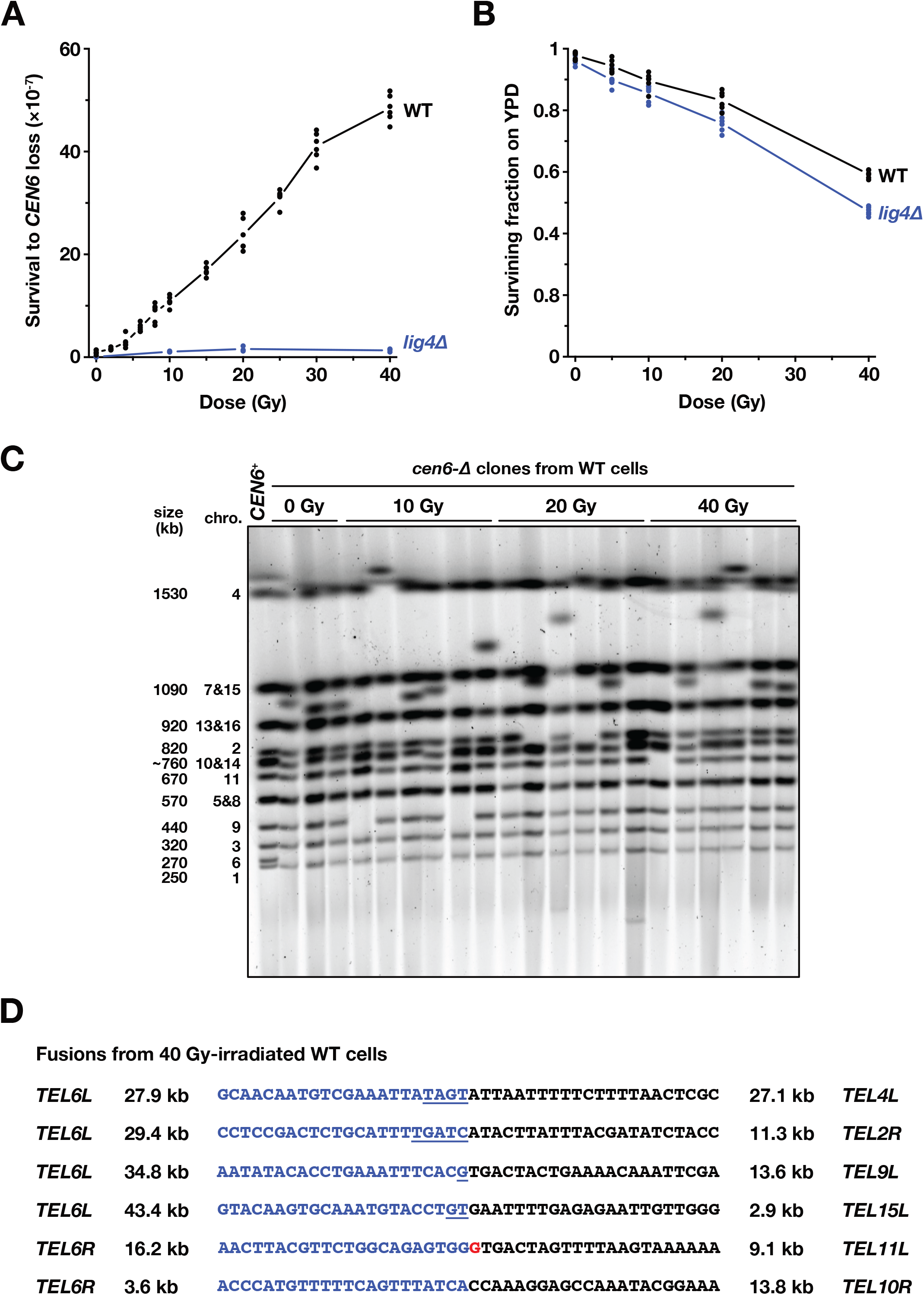
NHEJ-dependent chromosome fusions induced by ionising radiation. (**A**) Survival to *CEN6* loss in response to gamma ray irradiation (^137^Cs). (**B**) Cell survival in response to gamma ray irradiation. (**C**) Karyotype of *cen6Δ* clones from irradiated wild-type stationary cells. (**D**) Fusion point sequence in *cen6Δ* clones from irradiated wild-type stationary cells. Blue: chromosome 6 sequence. Underlined: Microhomology at the junction. Red: added non-homologous base. Distance from the closest telomere in kb.

Contrary to telomere-telomere fusions between chromosome 6 and another chromosome that always lead to viable outcomes once *CEN6* is deleted, we expect that the assay only selects a fraction of the chromosome 6 rearrangements stemming from radiation-induced DSBs. Fusions resulting in the loss of essential genes are not viable in haploid cells and therefore will not be selected. In practice, this severe constraint restricts viable chromosome fusions to the ones stemming from DSBs occurring within subtelomeric regions, which lack essential genes in yeast. On chromosome 6, this is 43,6 kb on the left end and 16.7 kb on the right end, representing together about 0.5% of the genome. On other chromosomes, non-essential subtelomeric regions are approximately 30 kb long and constitute together about 7.5% of the genome (combinatorial probability ~0.04%). In addition, we expect that the assay cannot select reciprocal translocations leading to unstable acentric chromosomes or interrupting essential genes. To test this prediction, we sequenced the genome of 6 clones with a radiation-induced chromosome fusion. All display a fusion between non-essential subtelomeric regions (**Fig. 4D**). The fusion points lack extensive homology, as expected for canonical NHEJ products. These data confirm that viability restricts the captured events to a small fraction of the radiation-induced chromosomal rearrangements. Spatial proximity between subtelomeres may also favour rearrangements between these regions (8, 50, 51). Despite these limitations, the assay offers a new, sensitive and quantitative method to address NHEJ repair of radiation-induced DSBs.

### Impact of dose fractionation on the frequency of radiation-induced rearrangements

First, to assess the specificity of radiation-induced rearrangements, we asked if photons of distinct energy have a similar ability to cause chromosomal rearrangements. In addition to 662 keV gamma rays, we irradiated cells with X rays of 68.5 and 20.4 keV effective energies respectively. All three lead to similar frequencies of chromosomal rearrangements (**Fig. 5A**). This result fits with previous observations in human cells and the predicted energies of the ionizing events (52). As expected, UVC irradiations, whose photon energy is too low to generate DSBs, do not lead to an increased survival to *CEN6* loss (**Fig. 5B**).

**Figure 5.**
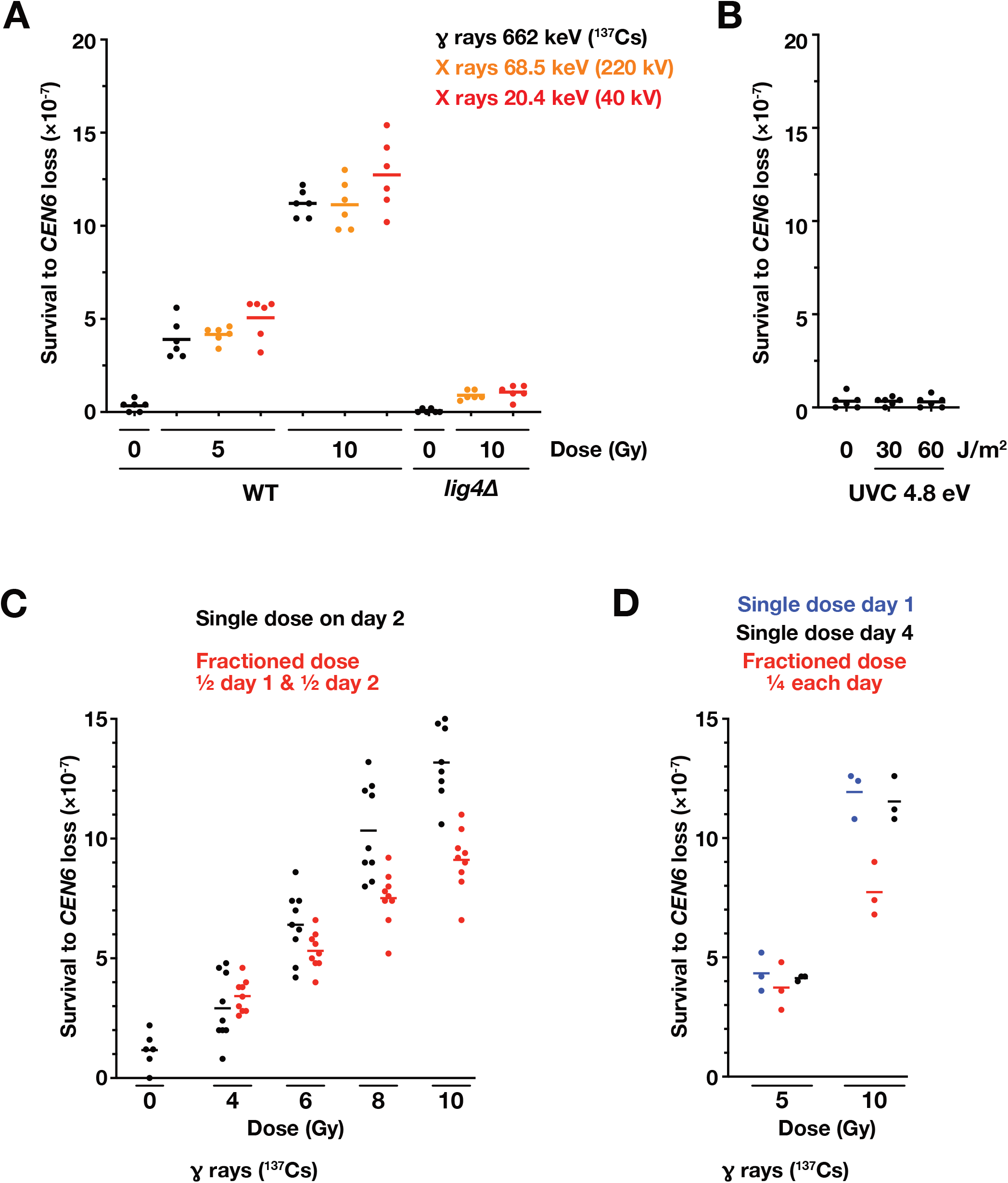
Impact of photon energy and dose fractionation on the frequency of radiation-induced rearrangements. (**A**) Survival to *CEN6* loss in response to X ray irradiations. (**B**) Survival to *CEN6* loss in response to UVC irradiations. (**C**) Survival to *CEN6* loss in response to dose fractionation. 24h interval between the two half-dose irradiations (^137^Cs). (**D**) Dose fractionated four times. 24h interval between the quarter-dose irradiations (^137^Cs).

NHEJ-dependent radiation-induced rearrangements result from the erroneous repair of co-occurring DSBs (16, 17, 53). This co-occurrence can stem from independent photon-matter interactions, each leading indirectly to the breakage of the two strands of distinct DNA molecules. In this scenario, the frequency of radiation-induced rearrangements is proportional to the square of the received dose and is sensitive to dose fractionation over time (which decreases the probability of co-occurrence). Alternatively, two co-occurring DSBs can stem from a single photon-matter interaction leading to the simultaneous breakage of two distinct DNA molecules (*i.e.* four strands). In this second scenario, the frequency of radiation-induced rearrangements progresses linearly with the received dose and is insensitive to dose fractionation over time.

To test the relevance of these scenarios in our assay, cells in stationary phase were irradiated either once with a full dose or twice with half-doses separated by a 24h interval. As shown in **Figure 5C**, fractioning a dose of 10 Gy (662 keV gamma rays) significantly reduces the frequency of captured rearrangements by about 30%. At lower doses, the impact of dose fractionation progressively lessens to disappear with an irradiation of 4 Gy. In addition, dose fractionation brings the frequency of the induced events closer to a linear (additive) increase with the dose (**Supplementary Fig. S3B**). To further assess the impact of dose fractionation, we exposed cells in stationary phase to doses of 5 and 10 Gy fractionated four times with a 24h interval between irradiations. Dose fractionation lowers the frequency of rearrangements induced by 10 Gy but does not significantly impact the frequency of events induced by a lower dose of 5 Gy (**Fig. 5D**). These results fit with a predominance of radiation-induced chromosomal rearrangements stemming from a single photon-matter interaction at lower doses (5 Gy or less). At higher doses (above 5 Gy), the rearrangements stemming from multiple independent photon-matter interactions likely predominate, explaining their sensitivity to fractionation.

In the previous experiment, 24h separate the fractionated doses. Next, we asked if shorter intervals also decrease the frequency of rearrangements induced by a dose of 10 Gy. As shown in **Figure 6A**, reducing the time between irradiations reduces the impact of dose fractionation. Consistent with this finding, a 10 Gy irradiation delivered over 14.5 hours at a low dose-rate of 0.0114 Gy/min generates less rearrangements than a 10 Gy irradiation delivered in a few minutes at a dose-rate of 2.6 Gy/min (**Fig. 6B**). These data suggest that DSBs created by the first irradiation remain capable of rearranging with DSBs created by the second irradiation for less than a few hours, consistent with the timescale of DNA repair by NHEJ. Once this delay has passed, chromosomal rearrangements induced by the second irradiation would only add up to the rearrangements induced by the first irradiation. Alternatively, can the lowering of rearrangement frequency in response to dose fractionation be a consequence of changes induced by the first irradiation, perhaps partially buffering cells against subsequent radiation-induced DNA damages? To assess the likelihood of this hypothesis, we tested the impact of a 10 Gy irradiation on cells previously exposed to 5 Gy (2.79 Gy/min, 24 hours between the two irradiations). As shown in **Figure 6C**, an initial 5 Gy irradiation does not lessen the frequency of rearrangements generated by a second irradiation of 10 Gy. This result suggests that yeast cells in stationary phase do not adapt to ionising radiation, at least regarding the occurrence of radiation-induced DSBs and their repair by NHEJ.

**Figure 6.**
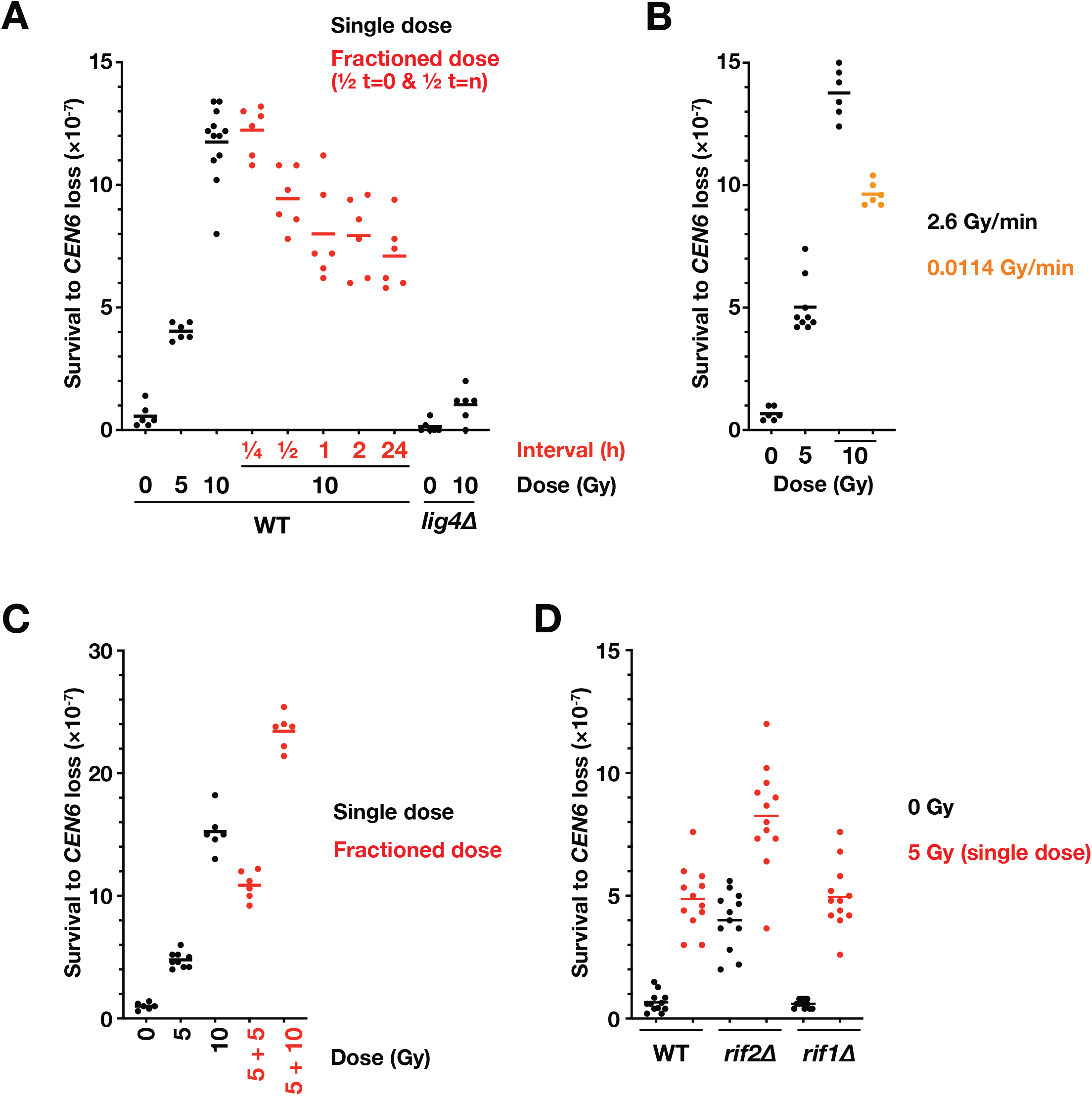
Impact of radiation density and telomere protection on the frequency of radiation-induced rearrangements. (**A**) Impact of the time between two irradiations of 5 Gy (^137^Cs). (**B**) Impact of a lower dose-rate (^137^Cs). (**C**) Impact of a first irradiation on the response to a second irradiation. 24h interval between the two irradiations (^137^Cs). (**D**) Impact of Rif2 and Rif1 loss (^137^Cs).

At lower doses (5 Gy or less), chromosomal rearrangements stemming from a single photon-matter interaction can explain the insensitivity to dose fractionation. One scenario is that one gamma ray sometimes causes indirectly the breakage of four DNA strands leading to two simultaneous DSBs. On the other hand, rearrangements at lower doses may occur between radiation-induced DSBs and double-strand ends of other origins. NHEJ-deficient haploid cells remain mostly viable in stationary phase (29, 49) (**Fig. 4B**), indicating that spontaneous DSBs of endogenous origin are rare events in these cells. DSB competence for NHEJ repair may also be short-lived, making them an unlikely source of ligatable double-strand ends. Another source could be telomere ends. If telomeres could sometimes fuse with radiation-induced DSB ends, changes in telomere protection efficiency should change the frequency of radiation-induced chromosomal rearrangements. To test this last scenario, stationary cells lacking Rif1 and Rif2 were irradiated. Rif1 loss, that reinforces telomere protection (**Fig. 3**), has no impact on the occurrence of radiation-induced rearrangements (**Fig. 6D**). As observed above (**Fig. 3**), Rif2 loss alone partially exposes telomeres to NHEJ, increasing the frequency of chromosome fusions. The rearrangements induced by a 5 Gy irradiation only add up to these events (**Fig. 6D**), indicating that Rif2 loss and ionising radiation act independently of each other. These results indicate that chromosomal rearrangements induced by 5 Gy do not usually occur between radiation-induced DSBs and telomeres. The more likely scenario is therefore that they are the erroneous repair products of two DSBs whose origin stems from a single photon-matter interaction.

## Discussion

In this work, we developed an assay to capture rare chromosome fusions. This tool shows telomere protection efficiency and measures the contribution of telomere dysfunction to genome instability in unchallenged normal cells. We found that telomere protection requires every pathway inhibiting NHEJ at telomeres and full telomerase activity. This new assay also captures chromosome rearrangements induced by ionising radiation and offers evidence for NHEJ-dependent chromosomal rearrangements stemming from single photon-matter interactions.

### Complementarity with the GCR assay

The chromosome fusion capture assay (CFC) presented here complements the widely-used gross chromosomal rearrangement assay (GCR) developed by Kolodner and colleagues (54–56). The latter detects rare losses of a chromosome fragment. Events selected by the GCR assay involve telomere addition at broken ends, interstitial deletions and chromosomal translocations. They rely mostly on error-prone mechanisms such as telomere *de novo* formation, microhomology-mediated end joining and break-induced replication. They still occur in NHEJ-deficient cells. By contrast, the CFC assay mostly captures events produced by the canonical Lig4-dependent NHEJ pathway. By giving us access to these events, it fills an important gap in the panel of tools available to study genome instability. The assay is amenable to systematic approaches and to genetic screens (e.g. (32)). In this work, we used it to address the efficiency of NHEJ inhibition at telomeres and the origin of the chromosomal rearrangements induced by ionising radiation.

### Low frequency of telomere fusions in wild-type cells

The CFC assay provides a first quantification of the basal rate of chromosome fusions occurring in unchallenged wild-type cells. In haploid *S. cerevisiae* cells reaching stationary phase, chromosome 6 is fused to another chromosome in about one every 10^7^ cells. With the assumption that all chromosomes are similar in this matter, the frequency of chromosome fusions would be about 10^−6^ per cell. Among the isolated fusions, we found a large fraction of telomere fusions. This indicates that telomere dysfunction in wild-type cells is a significant contributor to the basal level of chromosome fusions and therefore to time-depend mutagenesis in quiescence (57–59). Note that in budding yeast, an additional rescue pathway reverts telomere fusions into normal telomeres (34, 60, 61), limiting the impact of telomere fusions on genome stability.

The low frequency of telomere fusions observed in wild-type cells is the end product of the non-redundant actions of Rap1 and its two-cofactors Rif2 and Sir4. The CFC assay shows that Rif2 and Sir4 cannot fully back-up each other, explaining that both pathways are evolutionary stable. Efficient telomere protection also requires wild-type telomere length distribution since transient telomerase loss increases the frequency of fusions promptly. The absence of lag in the consequences of telomerase loss might be a hallmark of organisms like yeasts where telomeres are naturally short and telomerase constitutively expressed in all cells. By contrast, the absence of Rif1 diminishes the frequency of telomere fusions. This is a likely consequence of increased telomere length (and therefore increased Rap1 binding) and of reduced competition for Rif2 and Sir4 binding on Rap1. Then, what could be the evolutionary advantage of maintaining Rif1 at telomeres? Rif1 favours telomere stability during replication (42, 43, 62, 63) and this positive function may simply offset Rif1 negative impact on telomere protection against NHEJ.

### Evidence for NHEJ-dependent chromosomal rearrangements induced by single photon-matter interaction

The CFC assay provides a new way to quantify radiation-induced NHEJ-dependent chromosomal rearrangements. Its detection threshold is remarkably low, a few Gy. These are very low doses of ionising radiation for a small-genome organism such as budding yeast (1.4×10^7^ bp in a 2-3 μm^3^ nucleus (64)). CFC high sensitivity exposes radiation-induced chromosome fusions whose generation is linear (additive) with the dose and insensitive to radiation intensity. These events predominate at lower doses (≤5 Gy) and at lower dose-rates (≤1 Gy/h). Their linearity and insensitivity to intensity suggest that they stem from one photon-matter interaction (53). Since each of these rearrangements emanates from the repair of two DSBs, this argues that a single photon can lead to two DSBs.

**Figure 7** illustrates the molecular steps of this model. An incident gamma photon transfers a fraction of its energy to an electron as kinetic energy. Along its path through matter, the electron deposes this energy through multiple ionisation events (including the generation of secondary ionising electrons). Each energy deposition event can damage DNA directly but also indirectly by creating a transient and local high-concentration of reactive oxygen species (ROS) (17, 65, 66). If this small but highly reactive environment includes two DNA molecules in close proximity, a cluster of 4 single-strand breaks may form, leading to 2 co-incident DSBs. The 4 double-strand ends remain mobile relative to each other until they are captured by the NHEJ machinery whose first step is end synapsis (tethering) (9, 67). Breakage simultaneity, end proximity and rapid short-distance end diffusion result in frequent inaccurate synapsis (up to a 2/3 probability if pairing is random) and therefore inaccurate repair. In this model, chromosomal rearrangements are the products of rare but extremely mutagenic events, *i.e.* the breakage of two adjacent DNA molecules by a single track. An alternative model is that DSBs are produced in distinct energy deposition clusters along the electron path and would usually be distant at the time of their formation. This distance could favour accurate end synapsis and consequently accurate repair by NHEJ. It may therefore contribute less to mutagenesis. The relative likelihood of the two sequences of events remains to be determined.

**Figure 7.**
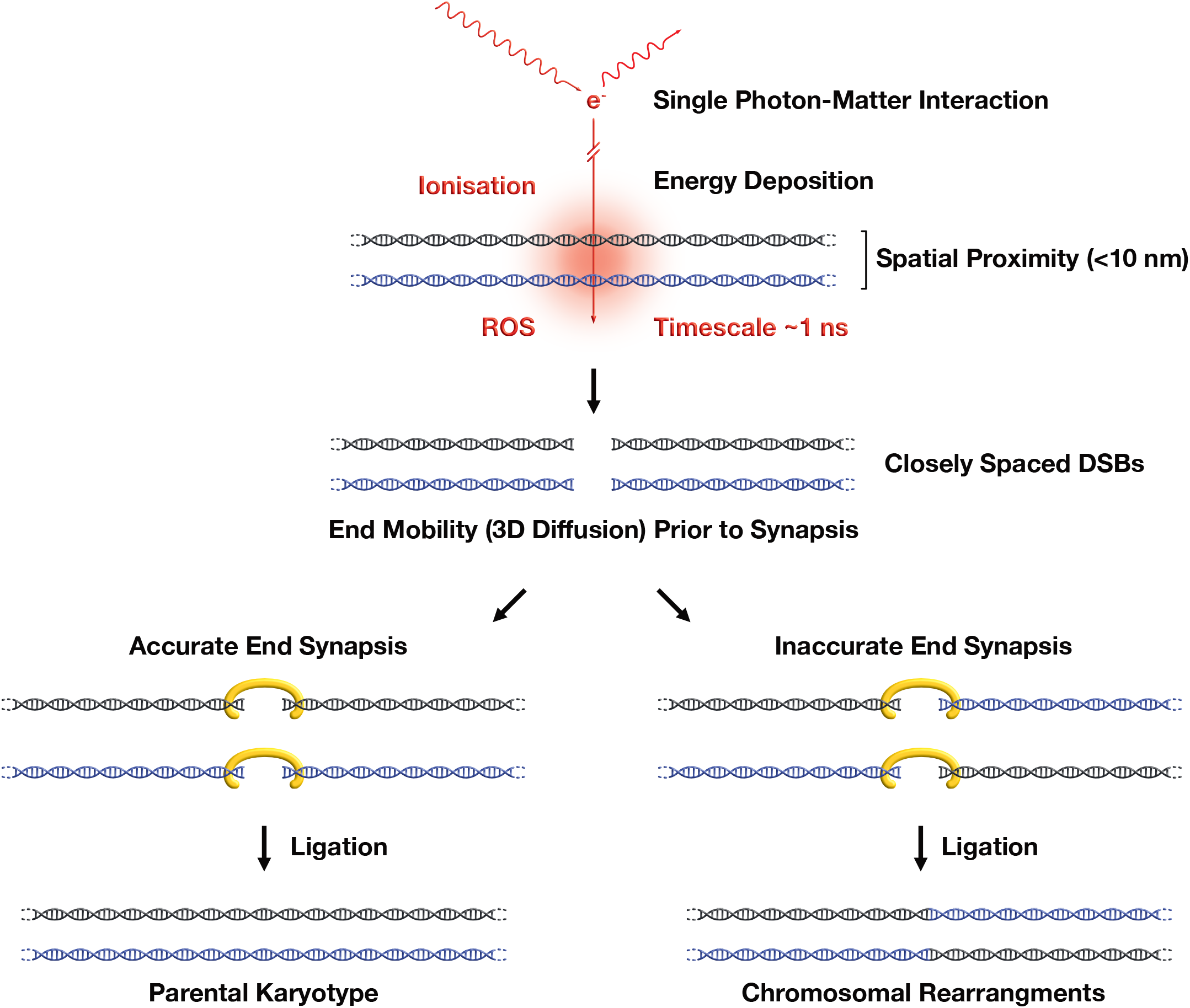
Model for NHEJ-dependent chromosomal rearrangements induced by single photon-matter interaction. Localised ionisation and ROS formation lead to two closely spaced DSBs and consequently frequent inaccurate end synapsis and repair by NHEJ.

The distinctive features of chromosomal rearrangements produced by single photon-matter interaction would be their linear (additive) increase with the dose and their insensitivity to radiation intensity. Exposure of mice to low dose-rates of ionising radiation causes an accumulation of chromosomal rearrangements that is remarkably linear with the dose (68, 69), in agreement with pioneering observations on plant cells (53). This suggests that the underlying mechanisms at the origin of these rearrangements are likely conserved in evolution from yeast to mammals. The CFC assay and the mechanism we propose provide a basis to address this issue at a molecular level. It will help to better assess the long-term impact of low-density irradiations and therefore the risks associated with medical radiography and the use of nuclear energy. In addition, the CFC assay can be a simple and sensitive tool to explore the significance of other putative causes of genome instability.

## Material & Methods

### Chromosome fusion capture and analysis

Strains, plasmids, and primers used in this study are listed in the attached spreadsheet. Telomeres and telomere fusions were amplified as previously described using primers TEL6R#31, X2, Y’2 and polyG14 (30, 31, 70).

To determine survival to *CEN6* loss, cells were grown to saturation in liquid rich medium (YPD) for 3 days at 30°C prior plating on galactose medium plates lacking leucine (5 × 10^7^ cells per plate or less if the frequency of survival to *CEN6* loss is above 4 × 10^−6^). Colonies were counted after 4-5 days at 30°C. A few strains used here lack the *klLEU2* marker inserted at *CEN6*. For those strains, cells were plated on galactose complete medium and the grown colonies were replicated on 5-FOA plates to identify clones having lost the *CEN6*-linked *klURA3* marker. On a sample of survivors (10 per plate), *CEN6* loss was confirmed by assessing the loss of a second *CEN6*-linked marker (*NAT^r^*).

### Sequencing and sequence analysis

Illumina paired-end whole genome sequencing files were processed with FastQC (v 0.11.4) for quality control and adapters sequences were removed with Cutadapt (v 1.13). Mapping was performed on the reference *S. cerevisiae* genome (Assembly sacCer3). Paired-ends from chromosome 6 with discordant mapping on the reference genome were identified with samtools (v 1.3.1) and a PERL script was used to align the reads together. Resolution was improved by visual inspection of the alignments on the breakpoint regions.

### Irradiations

Prior irradiation, dosimetry was performed with two different ionizing chambers as the recommendation of the AAPM’S TG-61 and depending on the irradiation configuration (71). We used a cylindrical ionizing chamber 31010 by PTW with a cavity of 0.125 cm^3^calibrated in ^137^Cesium air kerma free in air with the PTB reference facility number 1904442. This ionizing chamber was also calibrated in air kerma free in air in a range of 100-280 kV X-rays with the PTB reference facility number 1904444. In addition, we used a plane-parallel ionizing chamber with a cavity volume of 0.02 cm^3^calibrated in a range of 15-70 kV X-rays with the PTB reference facility number 1904440.

We performed gamma ray irradiations with a GSR D1 irradiator from the GSM Company. It is a self-shielded irradiator with four sources of ^137^Cs (a total activity around 180.28 TBq in March 2014). Cells in 6-well plates (1ml/well) were irradiated with an output of 2.6 or 0.0114 Gy/min.

We performed X ray irradiations with a SARRP (Small Animals Radiation Research Platform). Dosimetry protocol was calibrated to be adjusted to cells irradiation (72). Two configurations were used. The first configuration uses 0.8 mm of beryllium, 220 kV and 0.15 mm of copper for the additional filtration. The Half Value Layer (HVL) was measured at 0.677 mm of copper in order to obtain the spectrum and the effective energy of the photons: 68.5 KeV with SpeakCalc and XmuDAT AIEA’s software. The size of the field was 12*12 cm at FSD 35 cm and the dose output in these conditions was 2.7 Gy/min. The second configuration uses 0.8 mm beryllium, 40 kV and 1 mm of aluminium for the additional filtration. The HVL was measured at 0.886 mm of aluminium in order to obtain the spectrum and the effective energy of the photons: 20.4 KeV with the SpeakCalc and XmuDAT AIAE’s software. The size of the field was 15*11 cm at FSD 34 cm, the dose output in these conditions was 0.83 Gy/min. Due to the shape and the size of the field and to keep the homogeneity of the irradiation, only 4 wells on the 6 were irradiated (72).

## Aknowledgments

We thank Véronique Ménard for irradiation dosimetry, Didier Busso and Eléa Dizet (CIGEX platform) for plasmids, Clémentine Brocas and the CNRGH Sequencing facility for whole-genome sequencing, Claude Gazin, Karine Dubrana, Stefano Mattarocci, Florian Roisné-Hamelin, Zhou Xu, Gilles Fisher, Laurent Maloisel, Pablo Radicella, Paul-Henri Roméo, Gaëtan Gruel, Carmen Villagrasa, Nicolas Tang, Laure Sabatier and Christophe Carles for fruitful discussions and suggestions. This work was supported by funding from Fondation ARC, CEA Radiation biology programme, EDF and ANR (DICENs-ANR-14-CE10-0021-01).

## Supplementary Figures

**Supplementary Figure S1.**
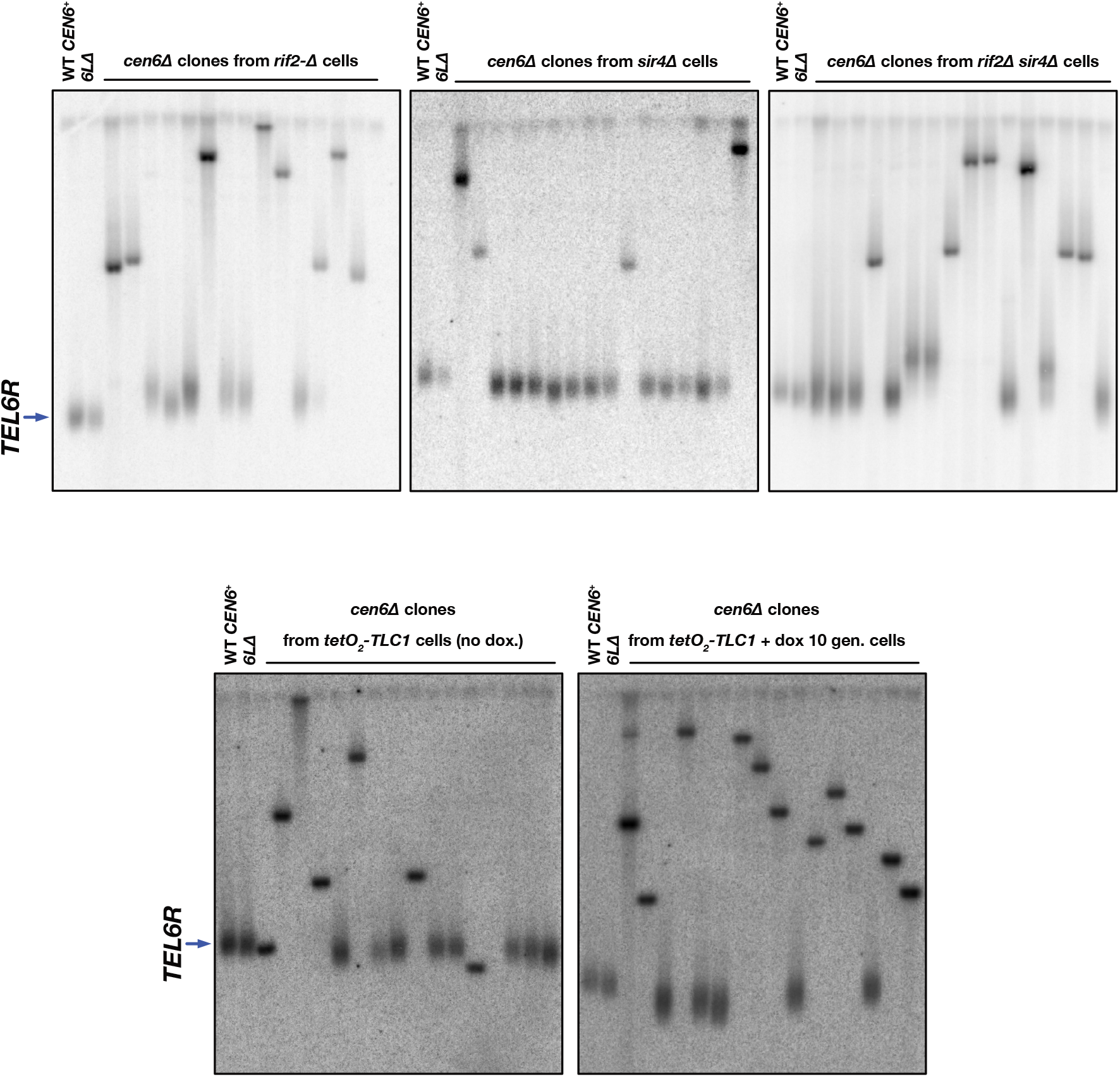
Telomere fusions at chromosome 6 right telomere among *cen6Δ* clones from stationary cells lacking Rif2, Sir4 or telomerase RNA template *TLC1*.

**Supplementary Figure S2.**
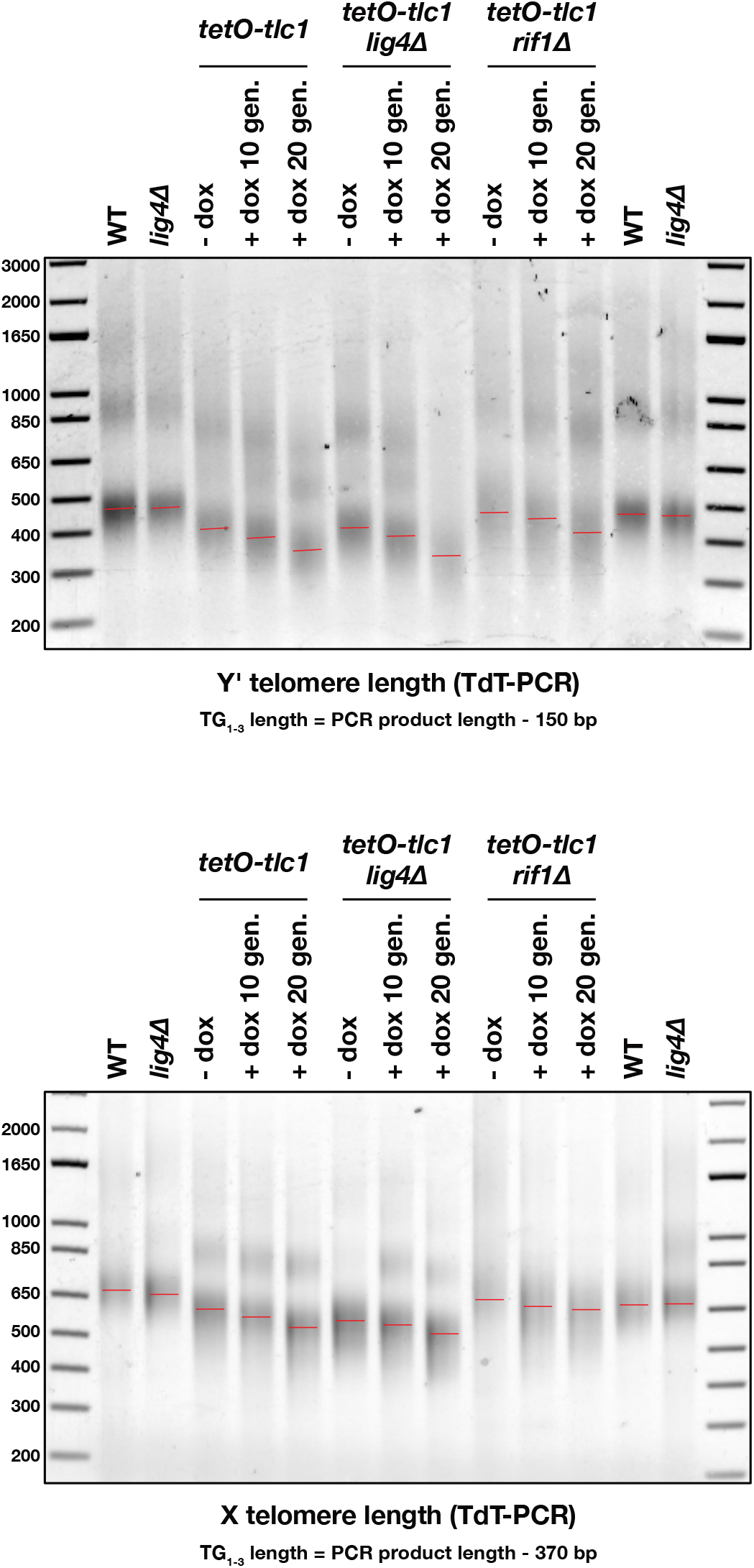
Telomere length in stationary cells lacking telomerase RNA template *TLC1* (TdT-PCR).

**Supplementary Figure S3.**
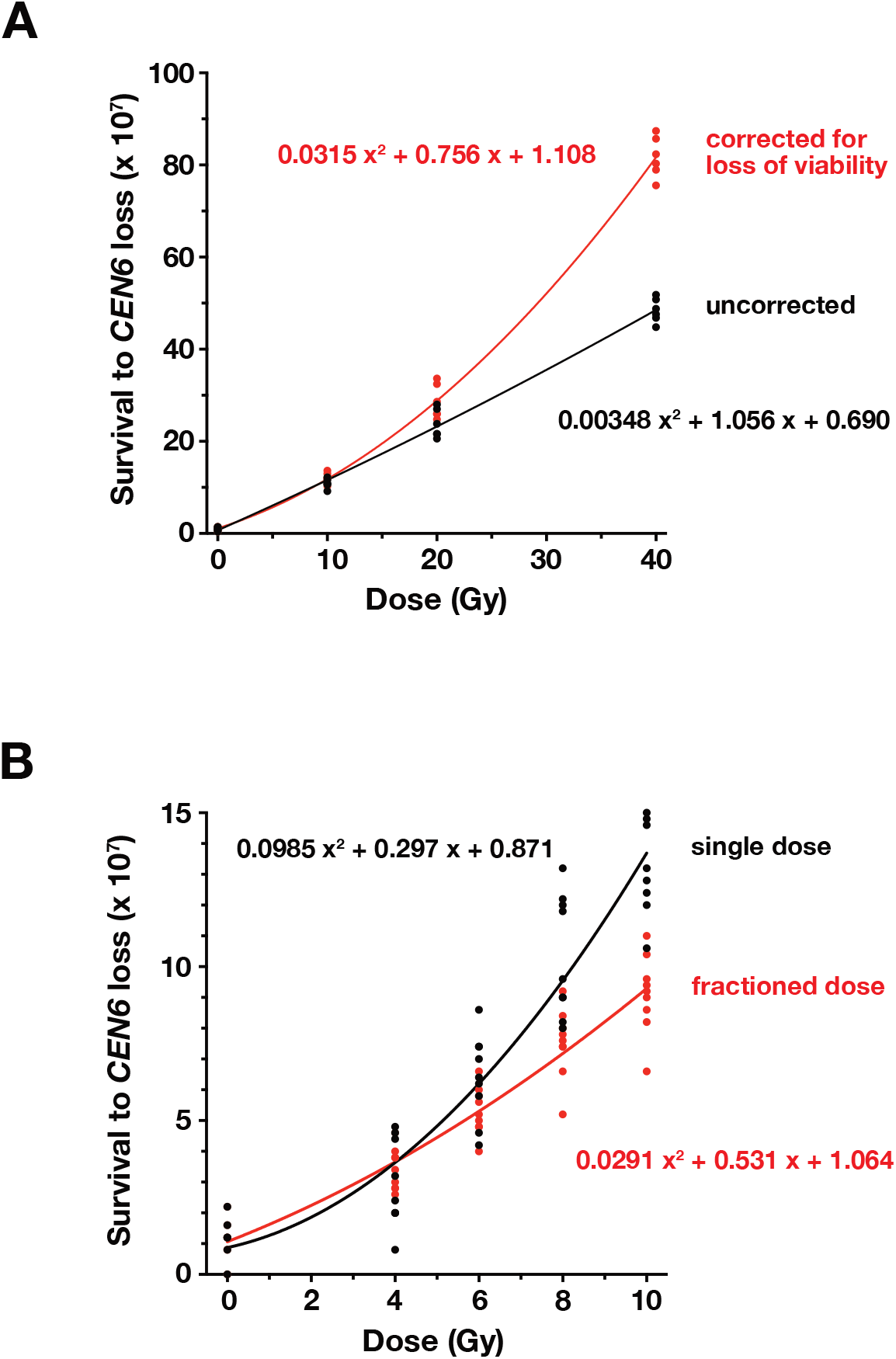
Quadratic regression of the relative survival to *CEN6* loss in response to ionizing radiation (^137^Cs). (**A**) Black curve: uncorrected. Red curve: corrected for loss of viability (each point divided by the mean surviving fraction on YPD). Data from Figure 4A&B. (**B**) Black curve: unfractionated doses. Red curve: fractionated doses. Data from Figure 5C.

